# Movement signaling in ventral pallidum and dopaminergic midbrain is gated by behavioral state in singing birds

**DOI:** 10.1101/2020.06.22.164814

**Authors:** Ruidong Chen, Vikram Gadagkar, Andrea C. Roeser, Pavel A. Puzerey, Jesse H. Goldberg

**Author notes:** The authors declare no competing financial interests.

## Abstract

Movement-related neuronal discharge in ventral tegmental area (VTA) and ventral pallidum (VP) is inconsistently observed across studies. One possibility is that some neurons are movement-related and others are not. Another possibility is that the precise behavioral conditions matter - that a single neuron can be movement related under certain behavioral states but not others. We recorded single VTA and VP neurons in birds transitioning between singing and non-singing states, while monitoring body movement with microdrive-mounted accelerometers. Many VP and VTA neurons exhibited body movement-locked activity exclusively when the bird was not singing. During singing, VP and VTA neurons could switch off their tuning to body movement and become instead precisely time-locked to specific song syllables. These changes in neuronal tuning occurred rapidly at state boundaries. Our findings show that movement-related activity in limbic circuits can be gated by behavioral context.

**Significance statement:** Neural signals in the limbic system have long been known to represent body movements as well as reward. Here we show that single neurons dramatically change their tuning to movements when a bird starts to sing.

## Introduction

Reward related signaling in ventral tegmental (VTA) and ventral pallidal (VP) regions is strongly state-dependent. For example, when an animal is hungry, a food predicting cue can drive dopamine release. But when sated, the DA system may be unresponsive (Ahn and Phillips 1999; Papageorgiou et al. 2016). Activity of VTA and VP neurons is also strongly driven by movements unrelated to reward [yin barter;engelhard;jin costa](Barter et al. 2015; Brooks 1986; Engelhard et al. 2019; Jin and Costa 2010). It remains unknown how these non-reward, movement related signals depend on what an animal is actually doing.

Zebra finches engage in ‘bouts’ of singing on and off during a typical day. Both during and outside these singing bouts, finches exhibit brief movements such as orienting their head and hopping from perch to perch (Eckmeier et al. 2008; Williams 2001). Recently, we and others discovered that VP and VTA neurons encode singing-related neural activity, including performance error signals important for song learning (Chen et al. 2019; Gadagkar et al. 2016; Hisey et al. 2018; Kearney et al. 2019; Xiao et al. 2018). These error signals functionally resembled reward prediction error signals observed in the limbic system (Humphries and Prescott 2010). We also discovered neurons with precisely time-locked firing to specific syllables in VP (Chen et al. 2019).

Here we investigate the movement-related firing properties of VP and VTA neurons and how they depend on whether birds are singing or not. In both VP and VTA, we discovered neurons that change their tuning to movement as birds transition from non-singing to singing states, demonstrating a gating mechanism for movement representation in limbic circuits commonly associated with reward and performance evaluation.

## Results

### Measuring neural activity and movement as birds transition into and out of singing states

To test if neural activity correlated with movement timing, we recorded movements with accelerometers attached to head-mounted microdrives (Fig. 1A). In the recording chamber most movements were head movements associated with orienting and whole body movements during hops. These transient movements occurred during both singing and non-singing periods in the day. Movements were more likely to occur right before onset of syllables (Fig 1B, peak movement onset probability 35 ± 2 ms before syllable onset, significant in 42/71 birds, assessed using bootstrap, methods), as previously reported (Gadagkar et al. 2016). At the level of singing bouts, movement probability peaked right before a bout of singing (Fig 1C, peak movement onset probability 38 ± 0.2 ms before bout onset, significant in 61/71 birds, bootstrap), and trended down after singing (Fig 1D, not significant). During non-singing periods, movements were longer on average and had higher variance (mean duration 106 ± 6 ms during singing vs. 159 ± 17 ms during non-singing, N=42418 movements during, 44850 outside singing, Fig 1E), consistent with more stereotyped movements during singing (Williams 2001). To control for these differences, we include only those movements that have similar duration and amplitude in subsequent analysis (N=35227 movements during singing and non-singing conditions, Fig 1E, methods).

**Figure 1.**
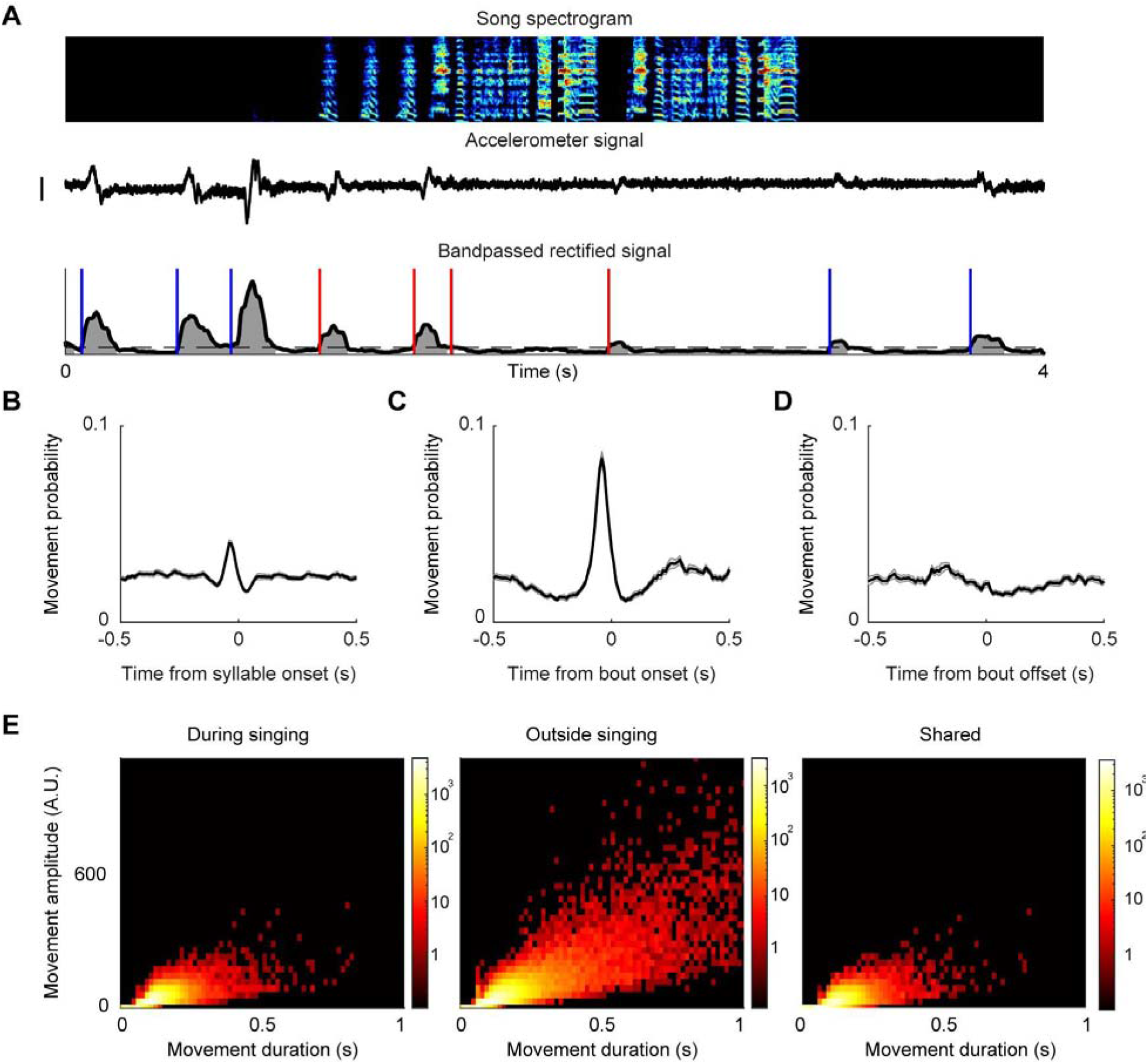
Measuring movement as birds transition into and out of singing states. A. (Top to bottom) song spectrogram, accelerometer signal, and bandpassed rectified accelerometer signal. Blue lines indicate onset of movements outside singing, red lines indicate onsets of movements during singing. Scale bar: 0.05mV. B. Average probability of movement onsets around syllable onsets (N = 71 birds). C. Same as C for song bout onsets. D. Same as C for song bout offsets. E. Left and middle: distribution of duration and amplitude of movements during and outside singing for all birds (n=71 birds, 42398 movements in singing, 138422 movements outside singing). Amplitude calculated as area under the curve of bandpassed rectified accelerometer signal (gray in A). Color axis: number of movements. Right: Same axes as left, with data from shared movements between both conditions (35227 movements from each condition).

### VP and VTA neurons encode movement timing

We recorded VP and VTA neurons as birds transitioned between singing and non-singing states (n=142 VP neurons, n=146 VTA neurons, n=71 birds) (Chen et al. 2019; Gadagkar et al. 2016). Many VP and VTA neurons exhibited activity that was precisely time-locked to movements outside of singing (26/142 neurons in VP, 77/146 neurons in VTA; rate extrema against randomly time-shuffled data with p<0.05, methods). Most neurons exhibited brief rate increases after movement onsets (latency to rate increase: 6.43 ± 3.8 ms, duration: 69.9 ± 3.1 ms, n=91/103 movement related neurons, Fig 2C), but neurons could also exhibit phasic decreases prior to movements (n=12/103 neurons).

**Figure 2.**
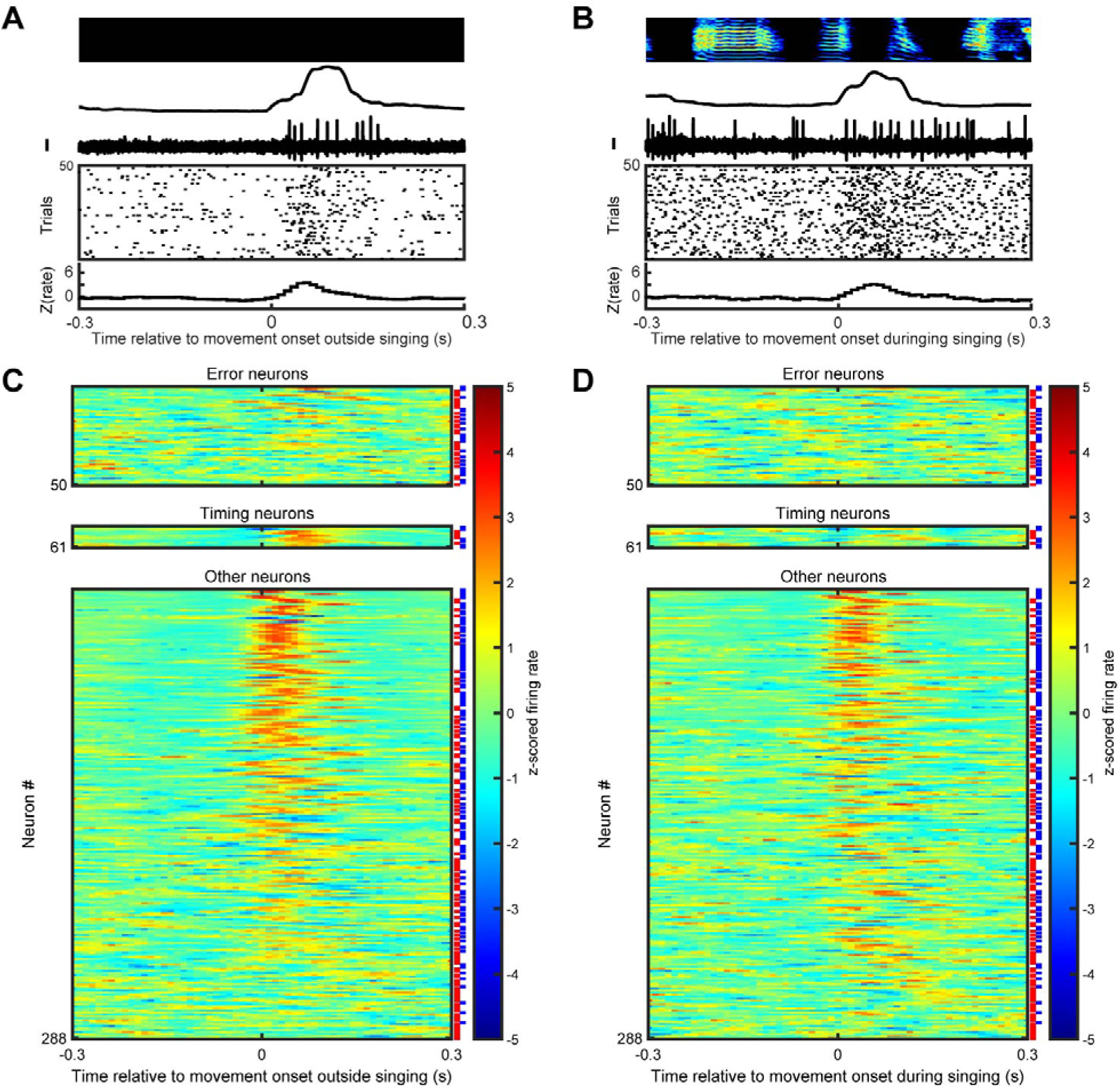
VP and VTA neurons exhibit movement-locked activity. A. Example VP neuron recorded outside singing. Top to bottom: spectrogram, bandpassed rectified accelerometer signal, neural voltage trace, raster plot of detected spikes, z scored firing rate. All plots are aligned to onsets of movements. Scale bar for neural activity is 0.25mV. B. Same neuron as in A but for movement onsets during singing. C. Z-scored firing rate aligned to movement onsets outside singing for all neurons, separated to error responsive, song timing related, and others. VP and VTA neurons are indicated by red and blue lines to the right of each row. Each group is sorted by maximum absolute response to movement onset. D. Same as C for movements during singing. Neurons are in the same order as in C.

During our recordings, we controlled perceived error with distorted auditory feedback (DAF) (Andalman and Fee 2009; Tumer and Brainard 2007). On randomly interleaved renditions of ‘target’ syllables, a brief, 50 millisecond snippet of sound was played through speakers surrounding the bird. In previous studies, we found that some VTA and VP neurons discharged differently to distorted and undistorted song renditions (Chen et al. 2019; Gadagkar et al. 2016). A small subset of these ‘error’ responsive neurons also exhibited movement-locked discharge (Fig 2C,D, n=1/23 VTA; n=4/27 VP error neurons were movement related, p<0.05, bootstrap, methods.). We also previously identified VP neurons with precisely timed song-locked discharge (operationally defined as intermotif pairwise correlation coefficient (IMCC) of 0.3 or higher (n=5 neurons in VP), methods). Interestingly, VTA neurons that were not error responsive (termed VTAother in (Gadagkar et al. 2016)) could also exhibit highly song-locked discharge (n=6/144 VTA neurons tested; 2/146 VTA neurons were recorded for less than 10 motifs and excluded from this analysis, see methods), reported here for the first time. These ‘song timing’ neurons could also be movement-locked outside of singing (Fig 2C,D, n=4/6 in VTA; n=3/5 in VP, p<0.05, bootstrap, methods).

### Movement-locked activity can depend on behavioral state

To test if movement responses changed during singing, we compared movement related changes in firing rate between singing and non singing states. Many neurons that were movement-locked outside of singing (Fig 2C) lost their tuning to similar movements during singing (Fig. 2D and Fig. 3, N=32/103 neurons with significant movement response only during non-singing states, including 5/5 movement-related error neurons and 4/7 movement-related timing neurons; N=71/103 neurons with significant responses in both singing and non-singing states).

**Figure 3.**
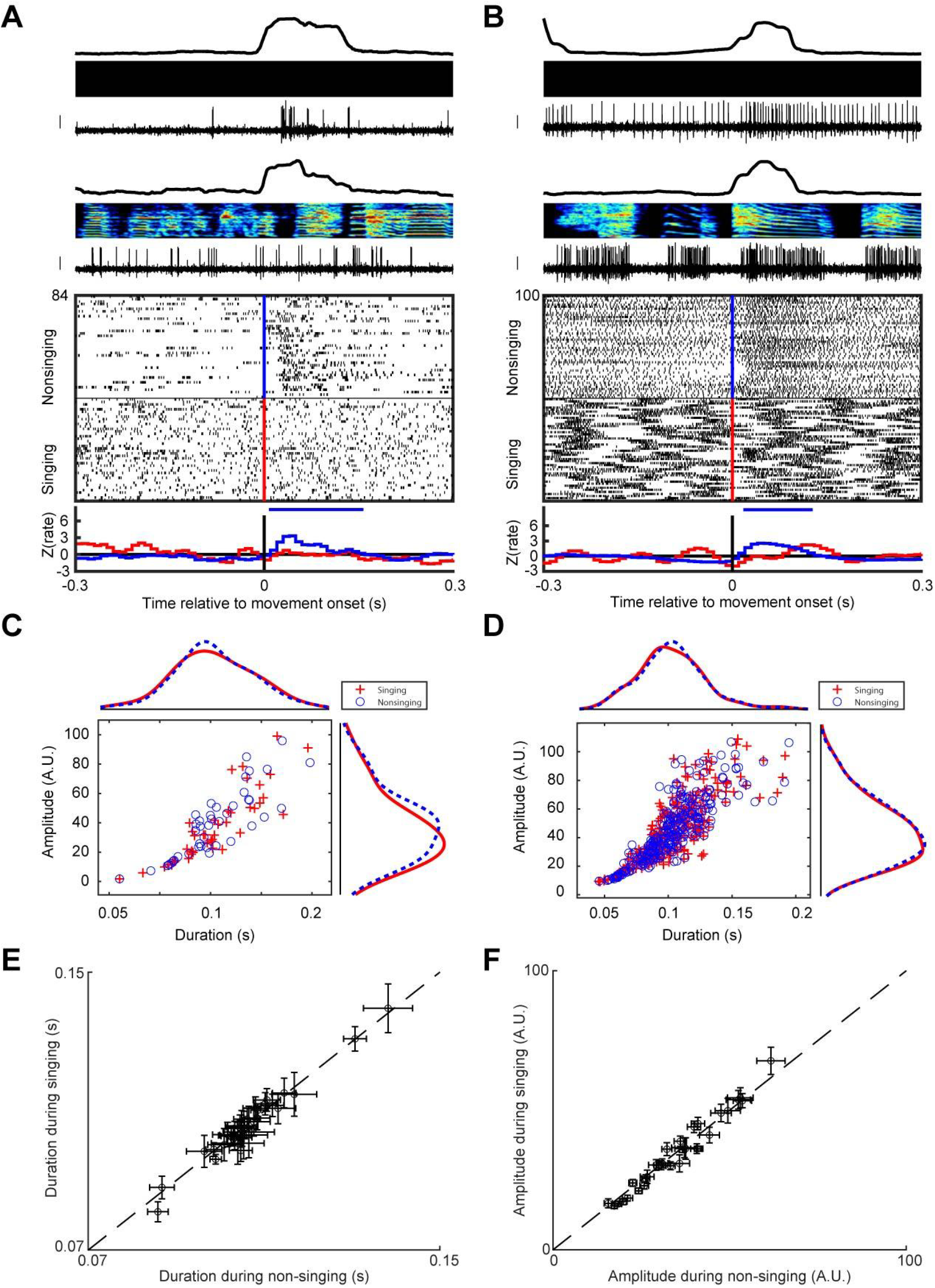
Example neurons that exhibit movement selectivity in the non-singing state only. A. (Top to bottom) bandpassed rectified accelerometer signal, spectrogram, voltage trace from an example VP neuron during non-singing and singing states, raster plot of spiking activity, z-scored firing rate histogram. Blue/red lines in the raster plot indicate movement onset during non-singing/singing. Horizontal bars indicate the duration of significant response to movement (z test, see methods). Scale bar for neural activity is 0.25mV. B. Same as A for an example VTA neuro. C. Duration and amplitude of movements shown in A for singing (red pluses) and non-singing (blue circles) states. Insets are kernel density plots of duration (top) and amplitude (right) of those movements. D. Same as C for the neuron in B. E. Duration of movements for state-dependent neurons. Circles are mean values from each neuron, error bars are S.E.M. F. Same as E for movement amplitude.

We wondered whether these state-dependent firing patterns could be attributed to differential distribution of movement types in singing vs. non singing states. We plotted the distribution of movement duration and amplitude from individual neurons in Fig 3B and Fig 3D and confirmed the movements were not systematically different in either duration or amplitude (two-sample Kolmogorov-Smirnov test, p>0.05, methods). As a group, state-dependent neurons shared similar movement feature distribution as other neurons (31/32 state-dependent neurons had similar feature distributions, K-S test, p>0.05. Mean +/-variance of movement duration and amplitude for each state-dependent neuron shown in Fig 3E,F).

### Movement related neurons switch to song-locked firing during singing

In both VP and VTA, movement related neurons could be precisely time locked to song syllables during singing (Fig. 4A,B, Chen et al. 2019). Because song timing and movement timing is correlated during singing (c.f. Fig 1B,C in ref (Gadagkar et al. 2016)), a purely movement locked neuron could show spurious correlation to song timing. To test this, we compared the magnitude of movement aligned rate modulation between singing and non-singing states. Movement modulation was quantified by the maximum in absolute value of z-scored rate histogram, and song-locked firing was quantified by intermotif cross correlation (Goldberg and Fee 2010; Kao et al. 2008; Olveczky et al. 2005). In previous study, we discovered that while many VP neurons exhibited discharge time-locked to song syllables, a specific subclass of bursting neurons, termed ‘song timing’ neurons, exhibited bursts that were extremely locked to sub-syllabic elements with millisecond timescale precision, and as a result exhibited high intermotif correlations of their spike trains (Methods). These neurons with significant time-locked firing to song showed higher movement selectivity outside singing than during singing (n=10/11 timing neurons with IMCC>0.3; n=47/71 song-locked neurons with significant IMCC. Fig 4C).

**Figure 4.**
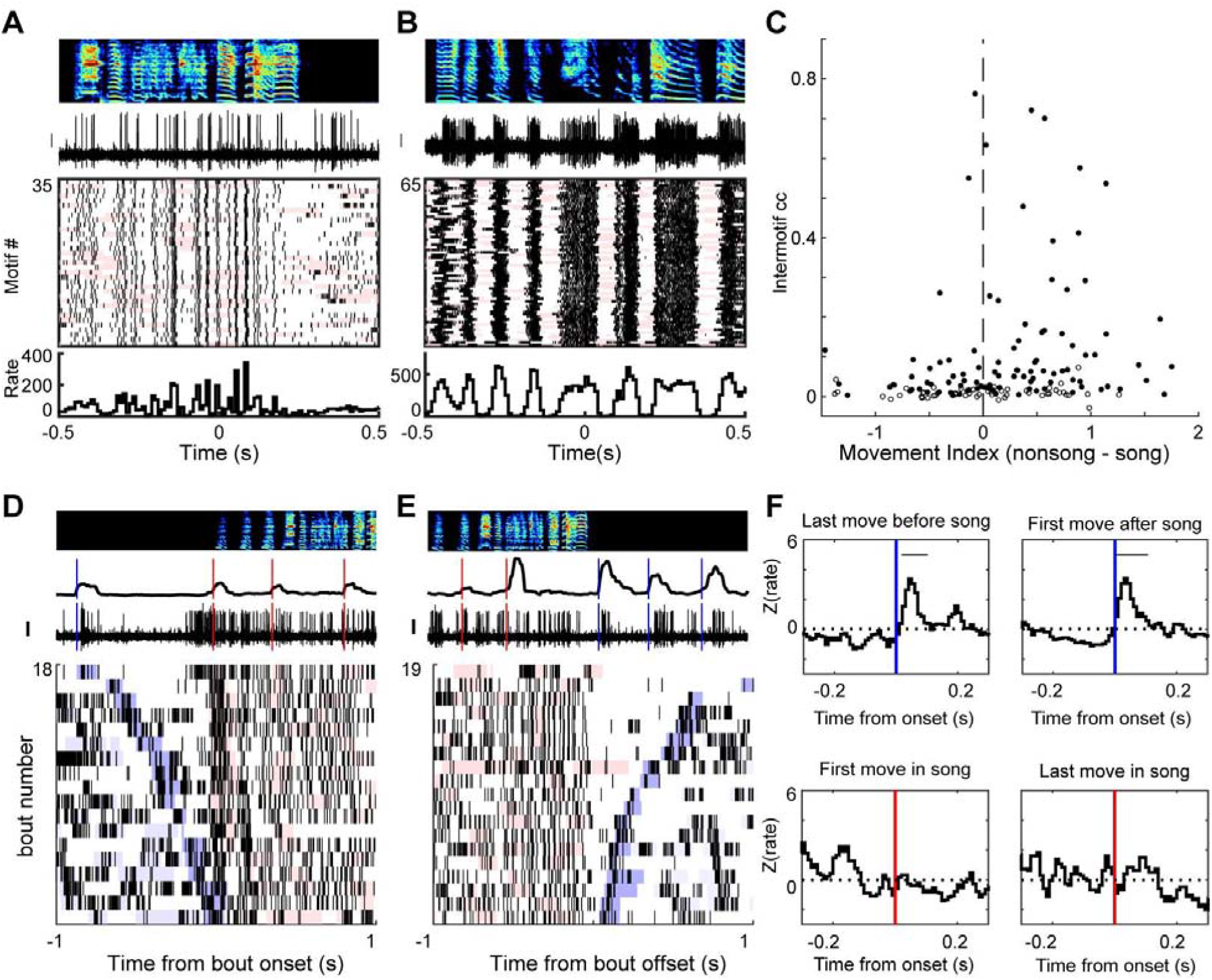
Example neurons that switch their tuning between song timing and movement at singing state boundaries. A. Movement related VP neuron time-locked to song. Same neuron as in Fig 3A. Top to bottom: song spectrogram, neural signal, rasters, and rate histogram aligned to song. Pink shades indicate movements during singing. Scale bar for neural activity is 0.25mV. B. Same as A for the VTA neuron in Fig 3B. C. Scatterplot of intermotif cross correlation and difference in movement index between non singing and singing states. Each dot is a movement related neuron. Filled circles are neurons with significant song-locked activity (p<0.05, methods). D. Example neural activity at transitions from non singing to singing, same neuron as A. Top to bottom: song spectrogram, bandpassed rectified accelerometer signal, neural signal, and rasters, aligned to onset of singing bouts. Blue and red lines indicate movement onsets outside and during singing. Blue shades in raster indicate movements outside song, with darker blue for the last movement before bout onset. Red shades indicate movements during singing. Raster sorted by the timing of the last movement before song. E. Same as D, but for transitions from singing to non singing. Raster sorted by timing of first movement after song. F. Top: z scored firing rate of neuron in D-E, aligned to movements immediately before (left) and after song (right). Bottom: same as top, for first and last movements during song. Blue/red bars indicate movement onsets outside/during song. Horizontal bars indicate significant response (p<0.05, z-test, methods).

### Movement signaling is rapidly gated at transitions to singing

To test if change in movement selectivity could occur immediately at state boundaries, we computed movement-aligned rate histograms for those movements surrounding state transitions (n=53 neurons with at least 10 transitions tested). Whereas the last movements occurring immediately before song bouts were correlated with bursts of firing, the first movements within a song bout were not (Fig 4D,F, 493 transitions, 17/53 neurons tested, Methods). Similarly, movements immediately following bout offsets were reliably associated with a burst of activity even though similar movements during singing were not (Fig 4E,F, 647 transitions, 14/53 neurons tested). Thus the change in tuning to movement can occur extremely rapidly (∼0.1 second timescale) at transitions to and from singing state.

## Discussion

By recording neural firing during singing and non-singing states in freely behaving birds, we discovered that movement-locked firing in VTA and VP neurons can be gated off during singing. In addition, neurons that were precisely time-locked to movements during non-singing became instead time-locked to song syllables (and not to body movements) during singing. These changes in tuning to distinct behavior modules could occur within tens of milliseconds at state boundaries. While basal ganglia and cerebellar neurons are known to be able to be differentially tuned to internally versus externally (e.g. cue-driven) movements (Strick et al. 1993; Van Donkelaar et al. 1999), to our knowledge such dramatic and rapid change in limbic tuning to movement across behavioral states has never been reported.

While the functional role of these state dependent movement representations is unclear, one possibility is that the context-dependent switch of representations between movement timing and song timing may reflect a common underlying algorithm that evaluates the quality of *any* motor program currently being produced, for example hop/orienting movements during non-singing versus syringeal movements during song. In this scenario, an outcome-weighted timing signal (such as a dopaminergic performance error signal) could be used to compute the predicted quality of ongoing performance, independent of the modality of the motor program.

Still, the switching of representation presents a puzzle for downstream neurons. For example, a neuron receiving input from a song timing related neuron during singing can reliably decode the time of song. However, as soon as singing stops, this same recipient neuron should also switch its decoding algorithm, for the incoming signal has changed. If this is the case, another input indicating the state change will be required. While the possible source of this proposed gating signal remains to be found, we note that previously discovered ‘SongOn’ and ‘SongOff’ neurons in VP (c.f. Fig. 4 in (Chen et al. 2019)) which turn on or off their activity during singing, act exactly like an ‘isSinging’ gate. The potential local connections between these cell types within VP is unknown.

One caveat in our study is that we were unable to fully distinguish between types of movements - e.g. hopping versus neck rotations. Instead, we have measured acceleration at the level of the head, and computed onset and offset timing of movements. While birds appear to move in similar ways during singing and non-singing periods (Yuan and Bottjer 2020), it’s possible that there are subtle systematic differences between ostensibly similar movements when performed during singing and non-singing. Future work with high speed video will be required to test this possibility. Notwithstanding, the cessation of reliable time-locked firing to movements during singing was striking.

Finally, in new analyses for the present paper, we discovered that non-error encoding VTA neurons (termed ‘VTAother’ in (Gadagkar et al. 2016)) could exhibit precise song-locked discharge (Fig 2C, 4B). Because error signals in the Area X projecting VTA neurons are temporally precise, timing signals in VTA could play a role in shaping dopaminergic signals important for song learning.

## Material and methods

### Subjects, surgery and histology

Subjects were 71 male zebra finches 74-355 days old singing undirected song. 61/71 birds and 269/288 neurons were new analyses of previously published datasets (Chen et al. 2019; Gadagkar et al. 2016). Animal care and experiments were approved by the Cornell Institutional Animal Care and Use Committee. All surgeries were performed with isoflurane anesthetization. Custom made microdrives carrying an accelerometer (Analog Devices AD22301), linear actuator (Faulhaber 0206 series micromotor) and homemade electrode arrays (5 electrodes, 3-5 MOhms, microprobes.com) were implanted into VP and VTA. VP implants (35 birds) were targeted using coordinates (4.4-5.4A, 1.1-1.5L, 3.5V, head angle 20 degrees). VTA implants (36 birds) were targeted using antidromic methods with stimulation in Area X (5.6A, 1.5L, 2.65V, head angle 20 degrees). After each experiment, small electrolytic lesions (30 μA for 60 s) were made with one of the recording electrodes. Brains were then fixed, cut into 100 μm thick sagittal sections for histological confirmation of stimulation electrode tracks and reference lesions.

### Singing and non-singing states

We separately analyzed neural activity and movement patterns in singing and non-singing states, and during transitions between these states, as previously described (Goldberg et al. 2010; Goldberg and Fee 2010). Bouts of singing was defined as consecutive syllables produced with gaps shorter than 300 ms. Non singing states were silent periods at least 300 ms away from syllables. In analysis of movement outside singing, only movements with onsets at least 300 ms away from song were included.

### Quantification of movement

An accelerometer (Analog Devices AD22301) was mounted on microdrives to detect gross body movements as described previously (Chen et al. 2019; Gadagkar et al. 2016). Briefly, movement onsets and offsets were determined by threshold crossings of the band-passed, rectified accelerometer signal. We further quantify the amplitude of each movement as the area under the curve of this signal (Fig 1A).

To assess the probability of movement near vocalizations, we computed the fraction of trials (period surrounding syllable onsets and bout on/offsets) in which movement onsets were detected in 10ms bins. To assess the significance of peaks in these probability functions, we compared the highest probability peak 1000 surrogate probability functions generated by randomly time-shifting movement onset relative to syllable onsets. Probability peaks exceeding the 95th percentile of surrogate probability maximum were considered significant.

To select similar movements shared between singing and non-singing states, we computed the joint distribution of duration and amplitude for all detected movements from each neuron, and restricted subsequent analysis to those movements that are shared in both conditions. In each bin, we randomly selected N movements from either state where N is the smaller number of movements in each state. Neurons that had at least 10 movements from each state were included for analysis. We confirmed the selected movements are sufficiently similar by performing two-sample Kolmogorov-Smirnov tests on the selected duration and amplitude distributions for each neuron.

### Analysis of neural activity

Neural signals were band-passed filtered (0.25-15 kHz) in homemade analog circuits and acquired at 40 kHz using custom Matlab software. Spike sorting was performed offline using custom Matlab software (courtesy Dmitriy Aronov). Firing rate histograms were constructed with 10 ms bins and smoothed with a 3-bin moving average.

To quantify movement locked neural response, we computed z-scored firing rates aligned to movement onsets during singing and non singing states using a 600 ms window centered on movement onsets as the baseline for each condition, respectively. This was chosen to reflect differences in degree of modulation rather than baseline firing rate between singing and non-singing conditions. Movement index was defined as the highest absolute z score within 100 ms before or after movement onsets. To assess the significance of these movement-locked rate modulations, we compared the highest rate peak and lowest nadir in movement onset-aligned rate histogram to 1000 surrogate rate histograms generated by randomly time-shifting spike trains. Rate peaks exceeding the 95th percentile of surrogate rate maximum and rate nadirs below the 5th percentile of surrogate rate minimum were considered significant.

To calculate the latencies and durations of movement responses, spiking activity within ±300 ms relative to movement onset was binned in a moving window of 10 ms with a step size of 5 ms. Each bin was tested against all the bins in the first 200 ms using a z-test. Response onset (latency) was defined as the first bin for which the next 4 consecutive bins were significantly different from the prior activity (z-test, P<0.05); response offset was defined as the first bin after response onset for which the next 2 consecutive bins did not differ from the prior (z-test, P>0.05). Response duration was the difference between the offset and the onset times.

To quantify movement-related responses at song state boundaries, we computed z-scored firing rates aligned to movement onsets using only movements either immediately before or after onsets and offsets of singing bouts. Those state dependent neurons that had at least 10 trials of each transition type were included in this analysis. Significant response to state boundaries was assessed with bootstrap method as above, and the duration of significant responses were quantified using z-test as above (Fig. 4F).

### Song timing related activity

Intermotif pairwise correlation coefficient (IMCC) was used to identify neurons that had highly time-locked firing to song motifs (timing neurons), as previously described (Chen et al 2019). Neurons with at least 10 motifs of singing were included in this analysis. Motif aligned instantaneous firing rate (IFR) was time warped to the median duration of all motifs, mean-subtracted, and smoothed with a Gaussian kernel of 20 ms SD, resulting in *r*_*i*_ for each motif. IMCC was defined as the mean value of all pairwise CC between *r*_*i*_ as follows:

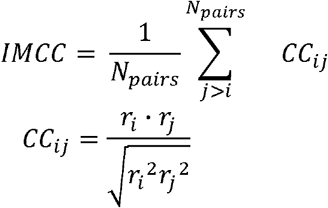

To assess the significance of IMCC values, we compared the true IMCC value to 1000 surrogate IMCC values generated by randomly time-shifting spike trains. IMCC values were considered significant if greater than the 95th percentile of the surrogate values.

### Error-related neurons

VP and VTA neurons were classified as error responsive (error neurons in Fig. 2) from previous studies (Chen et al. 2019; Gadagkar et al. 2016). Briefly, birds received syllable-targeted distorted auditory feedback (DAF) on randomly interleaved renditions. We compared target aligned activity between distorted and undistorted renditions, and those neurons with significant difference in firing following DAF were labeled as error neurons.

## Acknowledgement

This work was supported by funding from NIH R01NS094667.

## Notes

### Competing Interest Statement

The authors have declared no competing interest.

